# Genetic Rescue of a Lethal Wasting Mutation (*Syb1^lew/lew^*) by Neuron-Specific Expression of ECFP-Syb2

**DOI:** 10.64898/2026.05.22.727260

**Authors:** Yun Liu, Qiaohong Ye, Weichun Lin

## Abstract

Lethal-wasting (lew) is a spontaneous null mutation in the *Syb1/Vamp1* gene encoding the vesicular SNARE protein synaptobrevin 1 (SYB1/VAMP1). Homozygous *Syb1^lew/lew^* mice exhibit profound impairment in neuromuscular transmission and die around three weeks after birth. Pathogenic variants in human SYB1 are associated with a variety of disorders such as hereditary spastic ataxia and congenital myasthenic syndrome. Although Syb1 expression is highly enriched in neurons, it has also been reported in non-neuronal tissues, raising the possibility that non-neuronal defects contribute to the lethal phenotype. Here, we tested whether neuronal dysfunction is the primary cause of death in *Syb1^lew/lew^* mice and whether the closely related isoform synaptobrevin 2 (Syb2/VAMP2) can substitute for Syb1functin *in vivo*. We show that neuron-specific expression of *ECFP-Syb2* in *Syb1^lew/lew^* mice fully rescues lethality in *Syb1^lew/lew^*mice, restoring normal growth and motor function. Electrophysiological analyses demonstrate complete recovery of neuromuscular synaptic transmission, including spontaneous and evoked release as well as short-term plasticity. These findings establish that neuronal expression of ECFP-Syb2 is sufficient to prevent the lethal phenotype associated with Syb1 deficiency and demonstrate that Syb2 can functionally replace Syb1 at motor nerve terminals *in vivo*.

**Significance Statement:** Neuron-specific expression of Syb2 fully rescues survival, motor function, and neuromuscular transmission in *Syb1^lew/lew^* mice. These findings establish presynaptic Syb1 dysfunction as the primary cause of the lethal phenotype and demonstrate that Syb2 can fully substitute for Syb1 *in vivo*.

## Introduction

The mammalian nervous system expresses two major vesicular SNARE (soluble N-ethylmaleimide–sensitive factor attachment protein receptor) proteins, synaptobrevin 1(Syb1, VAMP1) and synaptobrevin 2 (Syb2, VAMP2), which mediate Ca²^+^-dependent synaptic vesicle fusion. Both Syb1 and Syb2 assemble with the target SNAREs syntaxin and SNAP-25 to form the ternary complex that drives synaptic vesicle fusion and rapid transmitter release (Jahn et al., 2024; Rothman and Orci, 1992; Sudhof, 2004; Sudhof and Rothman, 2009). Syb1 and Syb2 are highly homologous isoforms broadly expressed throughout the nervous system (Baumert et al., 1989; Elferink et al., 1989; Jacobsson et al., 1998; Li et al., 1996; Raptis et al., 2005; Sudhof et al., 1989; Trimble et al., 1988). Despite their similarity, disruption of either isoform results in neonatal lethality in mice (Schoch et al., 2001) or early postnatal lethality in mice (Liu et al., 2011; Liu et al., 2019; Nystuen et al., 2007), indicating essential and non-redundant roles in synaptic transmission and survival.

Syb1 exhibits a broad but selective distribution in the mammalian nervous system, with prominent enrichment in spinal and brainstem motor neurons and strong expression at the neuromuscular junction (NMJ) (Elferink et al., 1989; Jacobsson et al., 1998; Li et al., 1996; Liu et al., 2011). Consistent with this pattern, pathogenic SYB1 mutations are linked to human neuromuscular disorders, including hereditary spastic ataxia and presynaptic congenital myasthenic syndrome (Bourassa et al., 2012; Engel, 2018; Grewal et al., 2004; Meijer et al., 2002; Polavarapu et al., 2021; Rudolf et al., 2019; Salpietro et al., 2017; Shen et al., 2017; Yildirim et al., 2024). In mice, a null mutation in Syb1 causes the lethal-wasting (lew) phenotype and early postnatal death (Nystuen et al., 2007), accompanied by profound neuromuscular transmission defects, including reduced spontaneous and evoked release and impaired Ca²7 sensitivity and cooperativity (Liu et al., 2011).

In addition to its neuronal expression, Syb1 expression has also been reported in multiple non-neuronal tissues, including kidney, adrenal gland, liver, pancreas, thyroid, heart, and skeletal muscle (Ralston et al., 1994; Volchuk et al., 1994; Jo et al., 1995; Rossetto et al., 1996; Peters et al., 2006; Ferlito et al., 2010). This raises the possibility that the lethality of *Syb1^lew/lew^* mice may reflect defects in non-neuronal as well as neuronal cells. In addition, Syb1 and Syb2 are co-expressed at embryonic and juvenile NMJs (Liu et al., 2011; Liu et al., 2019), raising the question of whether these isoforms are functionally interchangeable at the NMJ.

To address these issues, we introduced neuron-specific expression of Syb2 to *Syb1^lew/lew^* mice by crossing them with transgenic mice expressing ECFP-tagged Syb2 under the Thy1 promoter (Thy1-ECFP-Syb2) (Liu et al., 2015). Remarkably, neuron-specific expression of ECFP-Syb2 fully rescued lethality, restored normal motor function, and normalized neuromuscular synaptic transmission in *Syb1^lew/lew^* mice. These findings demonstrate that Syb2 can functionally substitute for Syb1 at motor nerve terminals *in vivo*. Furthermore, because rescue was achieved through neuron-restricted expression, our results further indicate that Syb1 function in non-neuronal tissues is dispensable for survival. Together, this study supports the conclusion that Syb1 and Syb2 are largely functionally interchangeable at the NMJ and that neuronal synaptobrevin dosage is a critical determinant of viability in mice.

## Materials and methods

### Mice

The generation of *Thy1-ECFP-Syb2* transgenic mice was described in our previous studies (Liu et al., 2015). In these mice, the transgenic protein ECFP-Syb2 is expressed under the control of a neuron-specific Thy1 promoter. For simplicity, *Thy1-ECFP-Syb2* mice are hereafter referred to as *ECFP-Syb2* mice. *Syb1^+/lew^*heterozygous mice (C3H/HeDiSnJ-Vamp1^lew^/GrsrJ, stock number 004626) were obtained from the Jackson Laboratory (Bar Harbor, Maine, USA) and maintained on C57BL6 background. We bred *Syb1^+/lew^* mice with *ECFP-Syb2* transgenic mice to generate compound *Syb1^+/lew^;ECFP-Syb2* mice. These compound *Syb1^+/lew^;ECFP-Syb2* mice were subsequently crossed with *Syb1^+/lew^* mice to generate *Syb1^lew/lew^;ECFP-Syb2* mice.

The offspring were monitored daily for their external phenotypes and survival. Genotyping was carried out by PCR using the following primers. For *Syb1^lew/lew^* mice: wildtype allele, primer 1 - AGA GGG ACC AGA AGT TGT CAG and primer 2 - TGG CTG CTG GGT GTC TGA GG; mutant allele, primer 2 - TGG CTG CTG GGT GTC TGA GG and primer 3 - AGA GGG ACC AGA AGT TGT CAT. For *ECFP-Syb2* transgenic mice: forward primer-TGG TGA ACC GCA TCG AGC TG (from *ECFP*) and reverse primer - CGT TCA CCC TCA TGA TGT CC (from *Syb2*).

All experimental procedures were conducted in accordance with National Institutes of Health Guidelines and were approved by the University of Texas Southwestern Institutional Animal Care and Use Committee.

### Immunocytochemistry

Whole-mount muscle immunostaining was performed as previously described (Liu and Lin, 2024; Liu et al., 2008). Briefly, lumbrical or triangularis sterni muscles were fixed in 2% paraformaldehyde in 0.1 M phosphate buffer (pH 7.3) overnight at 4^°^C. Muscle samples were then incubated with Texas-Red conjugated α-bungarotoxin (α-bgt) (2 nM, Invitrogen, Carlsbad, California, USA) for 30 minutes, followed by incubation with primary antibodies diluted in antibody dilution buffer (500 mM NaCl, 0.01 M phosphate buffer, 3% BSA and 0.01% thimerosal) overnight at 4^°^C. The following primary antibodies were used: rabbit polyclonal anti-synaptotagmin2 (I735) (Pang et al., 2006), rabbit polyclonal anti-synaptobrevin1 (P938) and rabbit polyclonal anti-synaptobrevin2 (P939) (generous gifts from Dr. Thomas Südhof, Stanford University School of Medicine, Palo Alto, CA, USA). After extensive washes, muscle samples were incubated overnight at 4^°^C with Alexa Fluor 647-conjugated goat anti-rabbit IgG (1:600, Jackson ImmunoResearch Laboratories). Muscle samples were then washed with phosphate-buffered saline and mounted in Vectashield mounting medium.

To immunostain spinal cord sections, mice were anesthetized by isoflurane and transcardially perfused with 4% PFA. Spinal cords were then dissected and post-fixed in 4% PFA overnight at 4°C. Lumbar segments (L4–L6) were transversally sectioned at a thickness of 10 μm. Sections were labeled with TO-PRO-3 to visualize nuclei and immunostained by antibodies against synaptobrevin1 (P938, rabbit polyclononal) and synaptobrevin2 (CL69.1, mouse monoclonal). After washing, sections were incubated with Texas Red-Conjugated Goat Anti-Rabbit IgG (H+L) (1/600, Jackson ImmunoResearch Laboratories) and Alexa Fluor 488 goat anti-mouse IgG (H+L) (1:600, ThermoFisher). Sections were mounted in Vectashield mounting medium prior to imaging.

Fluorescent images were acquired using a Hamamatsu ORCA-285 camera or a Zeiss LSM 510 Meta confocal microscope.

### Electrophysiology

Experiments were performed on diaphragm muscles from P14 mice and lumbrical muscles from 5-month-old mice. Intracellular recordings were carried out as described previously (Liu et al., 2011; Liu et al., 2025). Briefly, diaphragm muscles with attached phrenic nerves or lumbrical muscles with attached plantar nerves were acutely isolated in oxygenated Ringer’s solution (136.8 mM NaCl, 5 mM KCl, 12 mM NaHCO_3_, 1 mM NaH_2_PO_4_, 1 mM MgCl_2_, 2 mM CaCl_2_, and 11 mM D-glucose; pH 7.3). End-plate regions were identified under a water-immersion objective (Olympus BX51WI) and impaled with glass micropipettes (20–40 MΩ resistance) filled with 2 M potassium citrate and 10 mM potassium chloride. Evoked end-plate potentials (EPPs) were elicited by suprathreshold stimulation (2–5 V, 0.1 ms) of the phrenic nerve using a suction electrode connected to an extracellular stimulator (SD9, Grass Telefactor, West Warwick, RI, USA). Muscle contractions were blocked by bath application of µ-conotoxin GIIIB (2 µM; Peptides International, Louisville, KY, USA) for 30 minutes prior to recording. Miniature end-plate potentials (mEPPs) and EPPs were recorded using an AxoClamp-2B intracellular amplifier and digitized with a Digidata 1322A interface (Molecular Devices, Sunnyvale, CA, USA). Data were analyzed with pClamp 9.0 (Molecular Devices) and Mini Analysis Program (Synaptosoft, Inc., Decatur, GA).

The amplitudes of mEPPs and EPPs were normalized to −75 mV by using the formula EPP_normalized_ = EPP × (−75/V_m_) where V_m_ was the measured resting membrane potential (Rozas et al., 2011). Quantal content (the number of acetylcholine quanta released in response to a single nerve impulse) was estimated using the direct method: dividing the mean normalized amplitude of EPPs by the mean normalized amplitude of mEPPs of the same cell (Boyd and Martin, 1956).

### Data analysis

Data are presented as the mean ± standard error of the mean (SEM). Kaplan-Meier survival curves were generated by using SigmaPlot 11.0. Statistical differences between control and mutant groups were determined by Student’s *t* test using Excel and were considered significant when the p value is less than 0.05.

## Results

### Syb1 and Syb2 show differential spinal cord distribution but are co-expressed at adult NMJs

Syb1 and Syb2 are the two major synaptobrevin isoforms in the mammalian nervous system and display distinct yet partially overlapping expression patterns. We previously showed that both isoforms are co-expressed in presynaptic terminals at the NMJ during embryonic and juvenile stages in mice (Liu et al., 2011; Liu et al., 2019). In adult mice, however, Syb1 has been reported to predominate at NMJs, with Syb2 expression markedly reduced or undetectable (Li et al., 1996; Takikawa and Nishimune, 2022).

To examine their distribution in the adult mice, we performed immunostaining on transverse lumbar spinal cord sections. Both Syb1 and Syb2 were detected throughout the gray matter (Fig. 1A), but with distinct regional enrichment: Syb1 was more prominent in the ventral horn, whereas Syb2 was enriched in the dorsal horn. Higher magnification revealed partial co-localization of the two isoforms within nerve terminals (Fig. 1B), consistent with previous findings in rats (Li et al., 1996).

**Figure 1.**
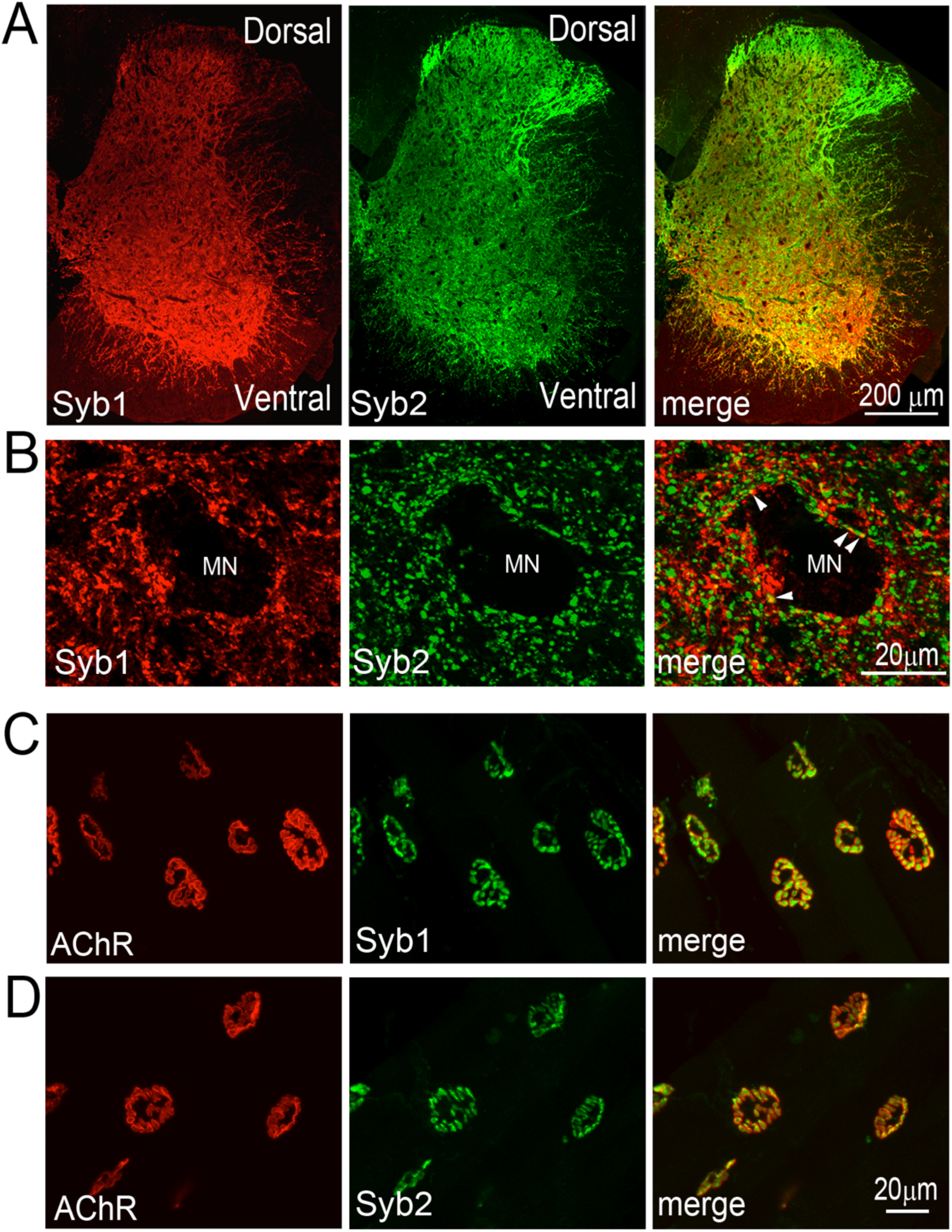
Expression of Syb1 and Syb2 in the spinal cords and the NMJ of adult wild-type mice. **A:** Transverse sections of lumbar spinal cord from a 4-month-old wild-type mouse were double labeled with antibodies against Syb1 (red) and Syb2 (green). Syb1 is enriched in the ventral horn, whereas Syb2 is more prominent in the dorsal horn. **B:** High-magnification images of the ventral horn region shown in A, highlighting a motor neuron (MN) and surrounding nerve terminals. Both Syb1 and Syb2 are detected in presynaptic terminals to the motor neuron. Syb1 and Syb2 are co-localized in a subset of terminals (arrowheads), with partial co-localization alongside terminals expressing either isoform alone. **C, D:** Whole-mount lumbrical muscles from 4-month-old wild-type mice were double-labeled with Texas Red-conjugated α-bgt and antibodies against Syb1(**C**) or Syb2 (**D**). Both Syb1 and Syb2 are detected at all motor nerve terminals of the NMJs.

We next assessed expression at adult NMJs using whole-mount lumbrical muscles. Double labeling with Texas Red–conjugated α-bungarotoxin and antibodies against Syb1 or Syb2 revealed that both isoforms are present at all labeled endplates (Fig. 1C, D), indicating that Syb1 and Syb2 remain co-expressed at adult NMJs.

### Neuron-specific ECFP-Syb2 expression does not perturb NMJ structure or function

ECFP-Syb2 transgenic mice express a neuron-specific fusion protein under the Thy1 promoter. In these mice, an exogenous ECFP-Syb2 fusion protein is expressed under the control of the Thy1 promoter, resulting in neuron-specific expression throughout the nervous system. (Liu et al., 2015). We examined distribution of ECFP-Syb2 in the adult spinal cord and at the NMJ. In transverse lumbar spinal cord sections from 5-month-old transgenic mice, CFP fluorescence was detected throughout the gray matter (Fig. 2A) and was enriched in nerve terminals surrounding motor neurons in the ventral horn (Fig. 2B). At NMJs, whole-mount triangularis sterni muscles were labeled with Texas Red–conjugated α-bungarotoxin to mark acetylcholine receptors and antibodies against synaptotagmin 2 to label presynaptic terminals. CFP fluorescence co-localized with synaptotagmin 2 at all labeled endplates (Fig. 2C, D), indicating that ECFP-Syb2 is targeted to presynaptic nerve terminals. NMJ morphology in transgenic mice was comparable to that of wild-type controls, suggesting that expression of the transgene does not disrupt synaptic structure.

**Figure 2.**
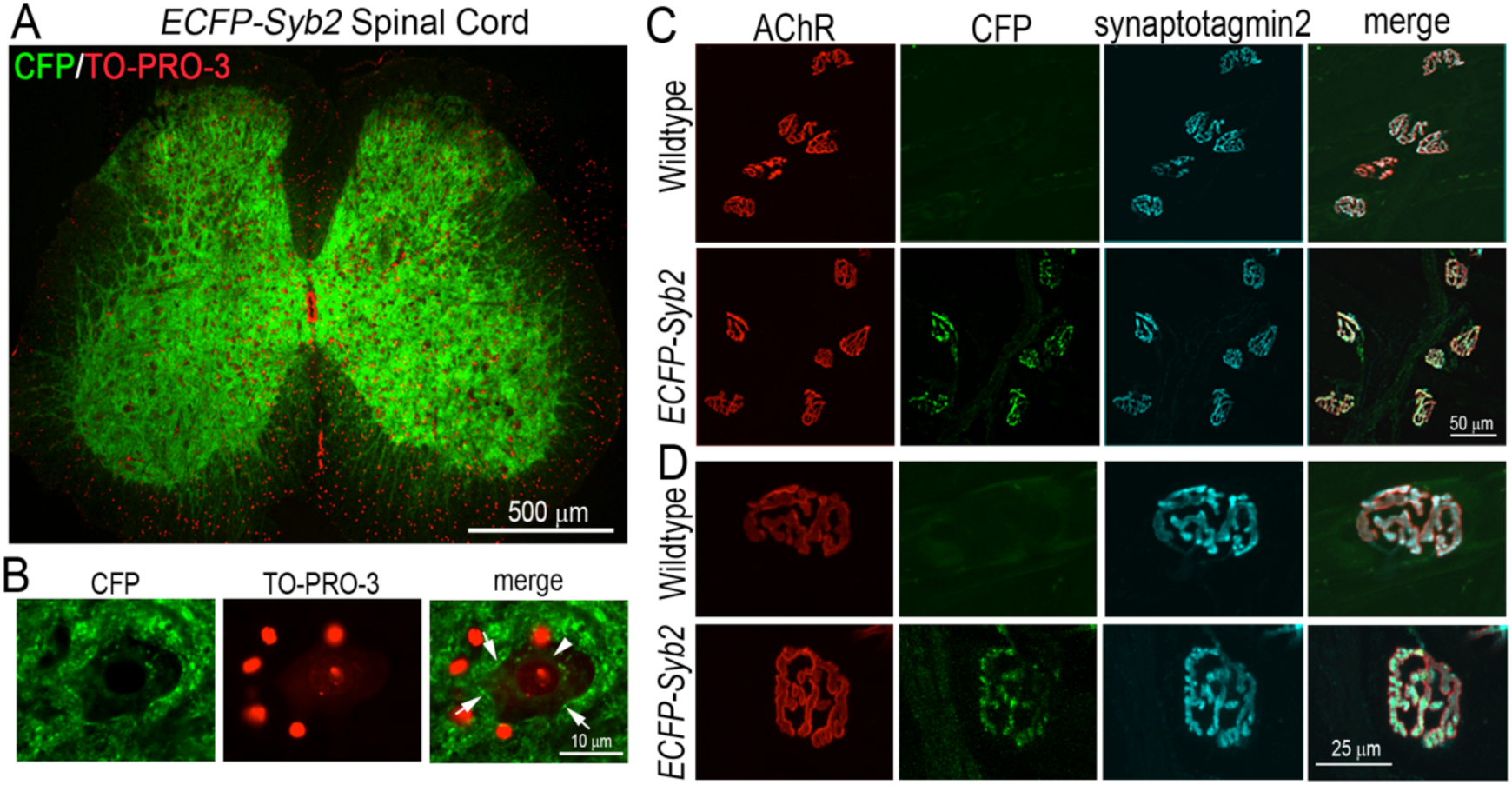
Expression of ECFP-Syb2 in spinal cord and the NMJ in ECFP-Syb2 mice. A: Transverse section of lumbar spinal cord from a 5-month-old *ECFP-Syb2* mouse labeled with TO-PRO-3 to visualize nuclei. CFP fluorescence is prominently detected in the gray matter. **B:** High-magnification image of a motor neuron in the ventral horn. CFP fluorescence is enriched in nerve terminals (arrows) contacting the motor neuron. The arrowhead indicates the nucleus of the motor neuron. **C:** Whole mounts of triangularis sterni muscles from 5-month-old wild-type and *ECFP-Syb2* mice were double-labeled with Texas Red-conjugated α-bungarotoxin (α-bgt) to label AChRs and antibodies against the presynaptic marker synaptotagmin 2 (Syt2) to label motor nerve terminals. ECFP-Syb2 is present at all NMJs and overlaps with the presynaptic marker Syt2. **D:** High-magnification images show that ECFP-Syb2 co-localizes with Syt2 at presynaptic terminals and is juxtaposed to AChR clusters, with NMJ morphology comparable to wild-type mice.

We next assessed whether expression of ECFP-Syb2 affects neuromuscular synaptic transmission. Electrophysiological recordings were performed on lumbrical muscles from 5-month-old transgenic and wild-type mice. Analysis of spontaneous neurotransmitter release showed no differences in miniature endplate potential (MEPP) frequency, amplitude, rise time, or half-width (Fig. 3). Similarly, evoked release was unaffected, with comparable endplate potential (EPP) amplitude, quantal content, rise time, and half-width between genotypes (Fig. 4A, B). During repetitive stimulation at 70 Hz, transgenic mice exhibited an EPP rundown profile indistinguishable from that of wildtype controls (Fig. 4C, D).

**Figure 3.**
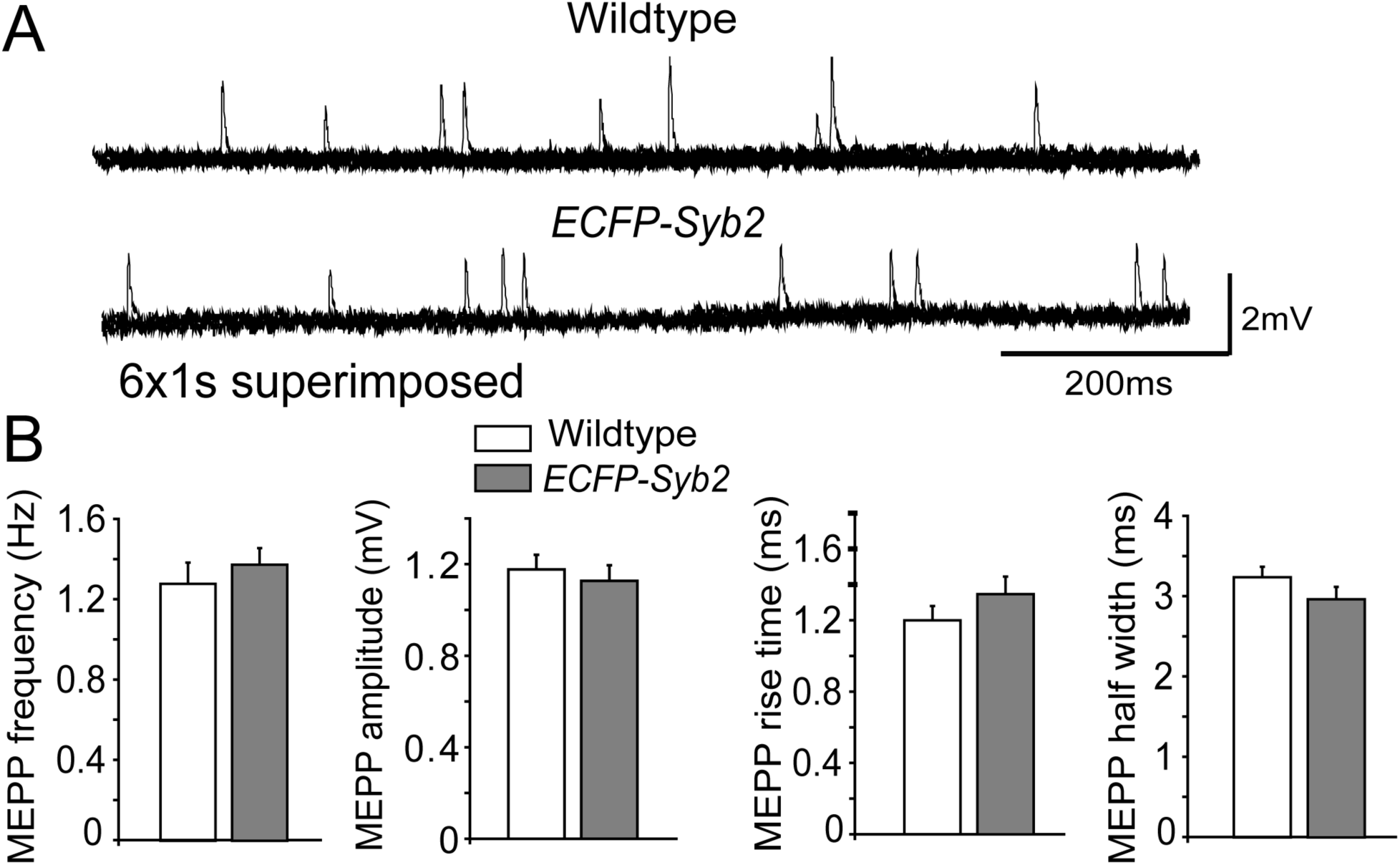
Spontaneous neurotransmission is normal at the NMJ in ECFP-Syb2 transgenic mice. **A**: Representative mEPP traces from wild-type and ECFP-Syb2 mice show comparable spontaneous activity. Sample mEPP traces represent 6 superimposed 1-second traces. **B**: Quantification of mEPP frequency, amplitude, risetime, and half-width reveals no significant differences between genotypes, indicating that ECFP-Syb2 expression does not affect baseline synaptic transmission. mEPP frequency : wild-type, 1.28 ± 0.11 Hz, *ECFP-Syb2*, 1.38 ± 0.08 Hz; MEPP amplitude: wild-type, 1.18 ± 0.06 mV, *ECFP-Syb2*, 1.13 ± 0.07 mV; rise time: wild-type, 1.2 ± 0.08 ms, *ECFP-Syb2*, 1.34 ± 0.09 ms; half width: wild-type, 3.25 ± 0.13 ms, *ECFP-Syb2*, 2.98 ± 0.15 ms. The number of mice and cells analyzed: wild-type: N = 3 mice, n = 32 cells; *ECFP-Syb2:* N = 3, n = 25.

**Figure 4.**
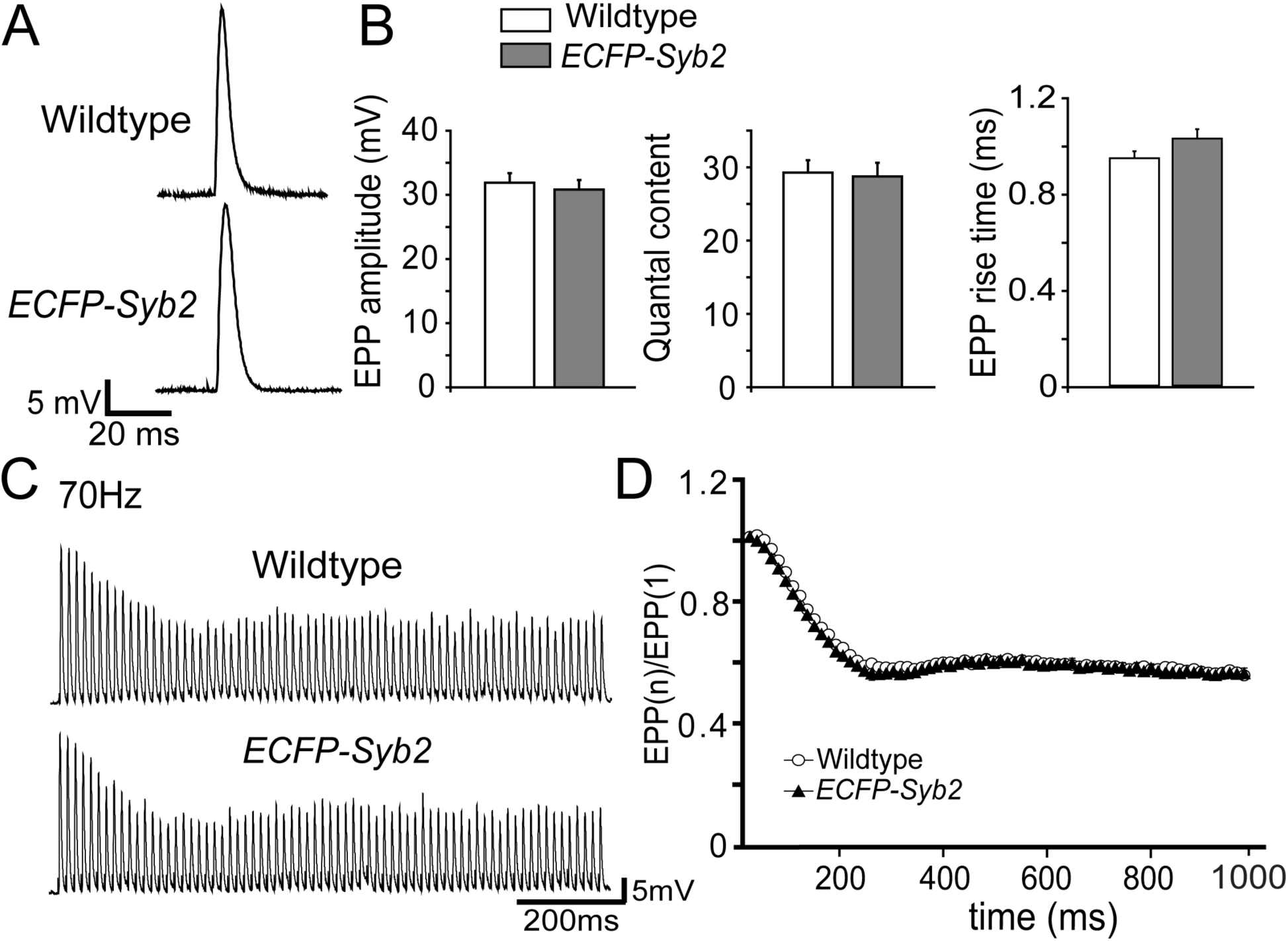
Evoked neurotransmission is normal at the NMJ in ECFP-Syb2 transgenic mice. **A**: Representative EPP traces from wild-type and ECFP-Syb2 mice show comparable evoked responses. **B**: EPP amplitude, quantal content, and rise time are unchanged in ECFP-Syb2 mice, indicating normal evoked release. No significant differences between wild-type and *ECFP-Syb2* mice. EPP amplitude: wild-type, 31.89 ± 1.49 mV, *ECFP-Syb2*, 30.83 ± 1.48 mV; quantal content: wild-type, 29.28 ± 1.68, *ECFP-Syb2*, 28.76 ± 1.85; rise time: wild-type, 0.94 ± 0.03 ms, *ECFP-Syb2*, 1.02 ± 0.06 ms. **C**: Sample EPP traces evoked by nerve stimulation at 70 Hz. **D**: Quantification of EPP rundown ratio in **E**. The rate of EPP run-down in *ECFP-Syb2* mice is similar with that in wild-type mice. The number of mice and cells analyzed are: wild-type: N = 3 mice, n = 30 cells; *ECFP-Syb2:* N = 3 mice, n = 21 cells.

Together, these results demonstrate that neuron-specific expression of ECFP-Syb2 does not alter NMJ formation, morphology, or synaptic function.

### Transgenic ECFP-Syb2 expression rescues lethal wasting (*lew*) and premature death in *Syb1^lew/lew^* mice

Because Syb1 and Syb2 are co-expressed at NMJs, we tested whether neuronal expression of ECFP-Syb2 could compensate for loss of Syb1 in *Syb1^lew/lew^*mice. To this end, *Syb1^+/lew^* mice were crossed with *ECFP-Syb2* transgenic mice to generate *Syb1^+/lew^*;*ECFP-Syb2* offspring, which were subsequently intercrossed to obtain *Syb1^lew/lew^*;*ECFP-Syb2* mice.

As previously reported, *Syb1^lew/lew^* mice exhibited severe growth retardation, progressive muscle weakness, and died prematurely (Liu et al., 2011; Nystuen et al., 2007). Among 55 offspring from 7 litters, 5 mice exhibited reduced body size and severe motor defects consistent with the reported phenotype of *Syb1^lew/lew^*mice. Genotyping confirmed that these 5 mice were *Syb1^lew/lew^*mutants lacking the ECFP-Syb2 transgene, and none survived beyond postnatal day 17 (P17). The remaining 50 mice appeared indistinguishable by gross inspection; genotyping identified 11 *Syb1^lew/lew^; ECFP-Syb2* mice within this group. Notebly, all *Syb1^lew/lew^; ECFP-Syb2* mice survived to adulthood, developed normally, and exhibited normal motor behavior (Fig. 5A).

**Figure 5.**
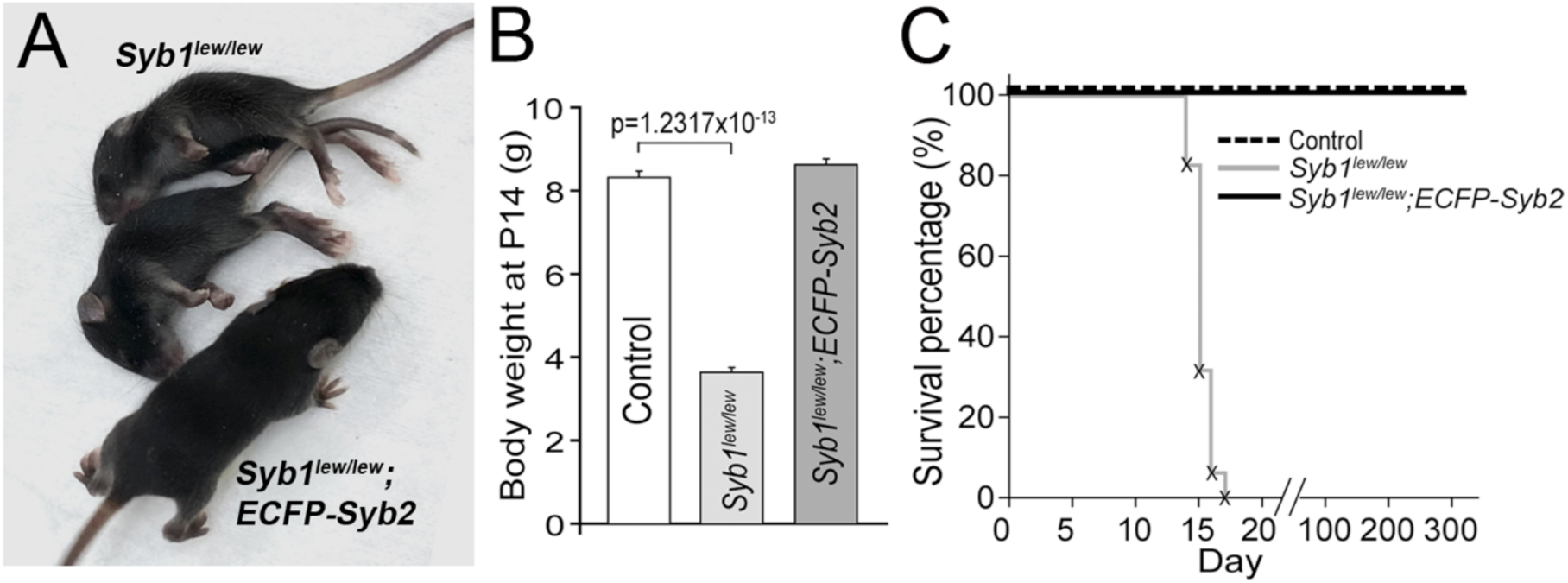
Transgenic ECFP-Syb2 expression rescues survival in *Syb1^lew/lew^*;ECFP-Syb2 mice. **A**: *Syb1^lew/lew^* mice (top two mice) display severe paralysis and immobility by P14. In contrast, their littermate *Syb1^lew/lew^;ECFP-Syb2* mouse is viable and exhibited normal motor behavior. **B**: Body weight is markedly reduced in *Syb1^lew/lew^* mice but restored to control levels in *Syb1^lew/lew^;ECFP-Syb2* mice. Body weight of *Syb1^lew/lew^* (3.64 ± 0.12 g, N = 5), *Syb1^lew/lew^;ECFP-Syb2* (8.62 ± 0.14 g, N = 11) and control (8.31 ± 0.15 g, N = 39, wild-type, *ECFP-Syb2* and *Syb1^lew/+^; ECFP-Syb2* mice are pooled together) mice at P14. **C:** Kaplan–Meier analysis shows complete lethality of *Syb1^lew/lew^*mice by P17, while ECFP-Syb2 expression fully rescues survival to control levels. Kaplan-Meier survival curves of control (N = 36), *Syb1^lew/lew^* (N = 16) and *Syb1^lew/lew^;ECFP-Syb2* mice (N = 33). *Syb1^lew/lew^;ECFP-Syb2* mice survived throughout the 310-day observation period. The “x” symbols on the survival curve for *Syb1^lew/lew^* mice indicate the time points at which deceased animals were identified.

Body weight was measured for all 55 mice at postnatal day 14 (P14). *Syb1^lew/lew^* mice showed a markedly reduced body weight (3.64 ± 0.12g, N = 5), whereas *Syb1^lew/lew^; ECFP-Syb2* mice had normal body weight (8.62 ± 0.14, N = 11), comparable to pooled controls (8.31 ± 0.15, N = 39; including wild-type, *ECFP-Syb2* and *Syb1^lew/+^;ECFP-Syb2* mice) (Fig. 5B).

Survival analysis further demonstrated complete rescue. We monitored the survival of *Syb1^lew/lew^;ECFP-Syb2* mice, *Syb1^lew/lew^* mice and control mice (pooled as above). All *Syb1^lew/lew^;ECFP-Syb2* mice (N = 33) survived throughout the 310-day observation period, indistinguishable from controls (n = 36) (Fig. 5C). In contrast, all *Syb1^lew/lew^*mice die by P17 (Fig. 5C).

Together, these results demonstrate that neuronal expression of ECFP-Syb2 fully rescues the lethal wasting phenotype and premature death caused by Syb1 deficiency.

### NMJ morphology is normal in *Syb1^lew/lew^*;*ECFP-Syb2* mice

Next we examined NMJ morphology in *Syb1^lew/lew^;ECFP-Syb2* mice. Whole-mount triangularis sterni muscles from P14 control, *Syb1^lew/lew^* and *Syb1^lew/lew^;ECFP-Syb2* mice were labeled with Texas Red–conjugated α-bungarotoxin and antibodies against Syb1. As expected, Syb1 immunoreactivity was detected at control NMJs but was absent in both *Syb1^lew/lew^*and *Syb1^lew/lew^; ECFP-Syb2* mice. In contrrast, GFP fluorescence from the ECFP-Syb2 transgene was observed exclusively at NMJs in *Syb1^lew/lew^;ECFP-Syb2* mice (Fig. 6A), confirming targeted expression at presynaptic terminals.

**Figure 6.**
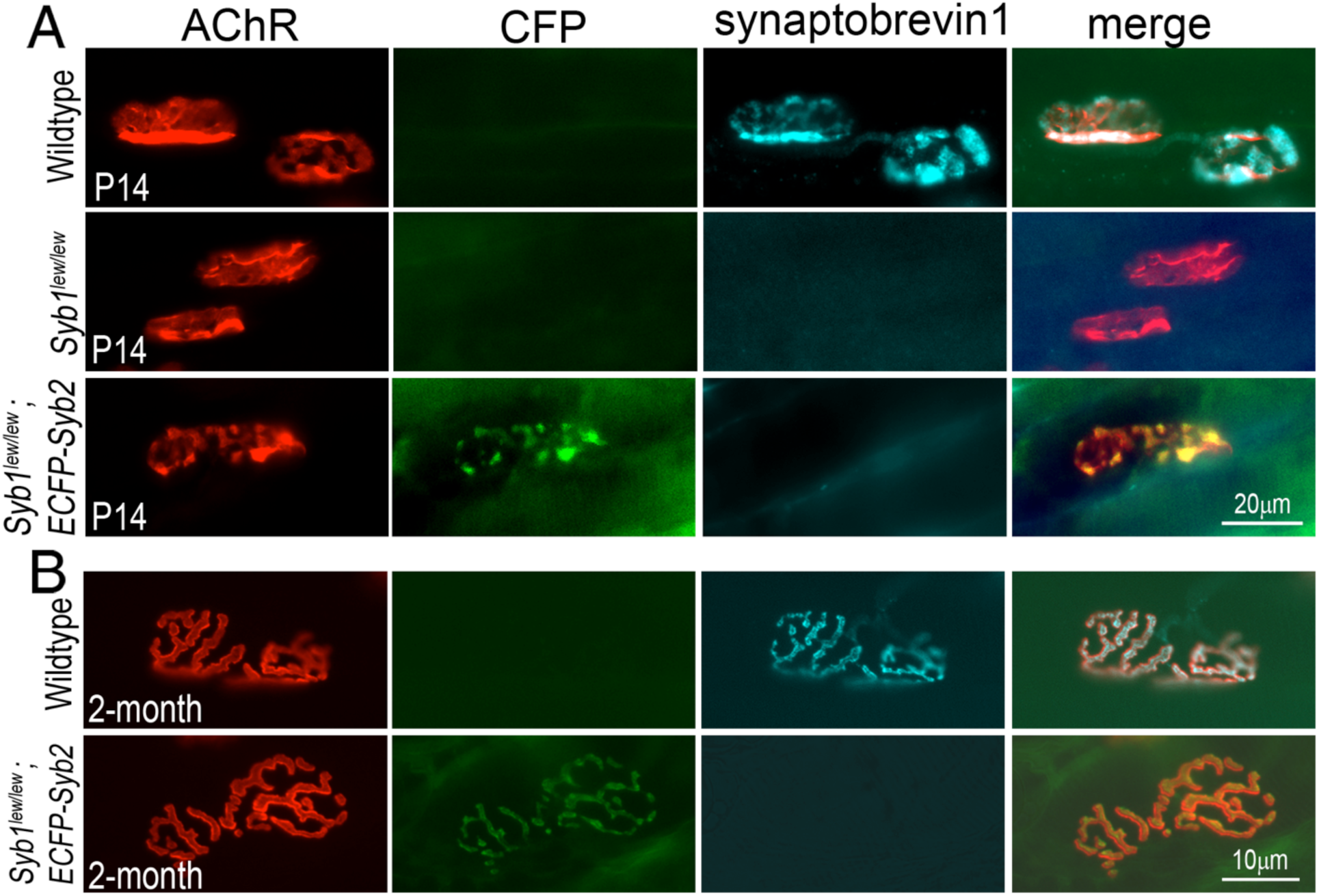
Normal morphology at the NMJ in *Syb1^lew/lew^;ECFP-Syb2* mice. **A**: Whole mounts of triangularis sterni muscles from wild-type, *Syb1^lew/lew^* and *Syb1^lew/lew^;ECFP-Syb2* mice at P14 were double-stained with Texas Red-conjugated α-bgt and antibodies against Syb1. Syb1 is detected at wild-type NMJs but absent from either *Syb1^lew/lew^* or *Syb1^lew/lew^;ECFP-Syb2* NMJs. ECFP-Syb2 is detected at NMJs of *Syb1^lew/lew^;ECFP-Syb2* mice. **B**: Whole mounts of triangularis sterni muscles from wild-type and *Syb1^lew/lew^;GFP--Syb2* mice at 2 months of age. NMJ morphology in S*yb1^lew/lew^;ECFP-Syb2* mice is comparable to that of wild-type NMJs.

To assess adult NMJs, we analyzed NMJ morphology at 2 months of age. In *Syb1^lew/lew^;ECFP-Syb2* mice, NMJs exhibited normal maturation and were stably maintained, with morphology comparable to that of control NMJs (Fig. 6B).

### Transgenic ECFP-Syb2 expression fully restores neuromuscular transmission in *Syb1^lew/lew^* **mice**

As reported previously, *Syb1^lew/lew^* mice exhibit severely impaired neurotransmitter release at NMJs, characterized by reduced spontaneous and evoked synaptic activity and enhanced short-term facilitation (Liu et al., 2011). Because the *ECFP-Syb2* transgene rescues the survival and motor deficits of *Syb1^lew/lew^*mice, we next tested whether it also restores neurotransmission at Syb1-deficient NMJs. To address this, we performed electrophysiological analyses of diaphragm muscles from control, *Syb1^lew/lew^*, and *Syb1^lew/lew^;ECFP-Syb2* mice at P14.

Spontaneous release was first assessed by measuring miniature endplate potentials (MEPPs). MEPP frequency was markedly reduced in *Syb1^lew/lew^* mice (3.29 ± 0.38 event/minute) compared to controls (10.21 ± 0.86 events/min). In contrast, MEPP frequency was robustly restored in *Syb1^lew/lew^;ECFP-Syb2* mice (11.06 ± 0.91 event/minute), comparable to control (10.21 ± 0.86, event/minute) (Fig. 7). MEPP amplitude was unchanged across genotypes (control: 2.1 ± 0.09 mV; *Syb1^lew/lew^*: 1.98 ± 0.14 mV; *Syb1^lew/lew^;ECFP-Syb2*: 2.04 ± 0.07 mV). (Fig. 7C).

**Figure 7.**
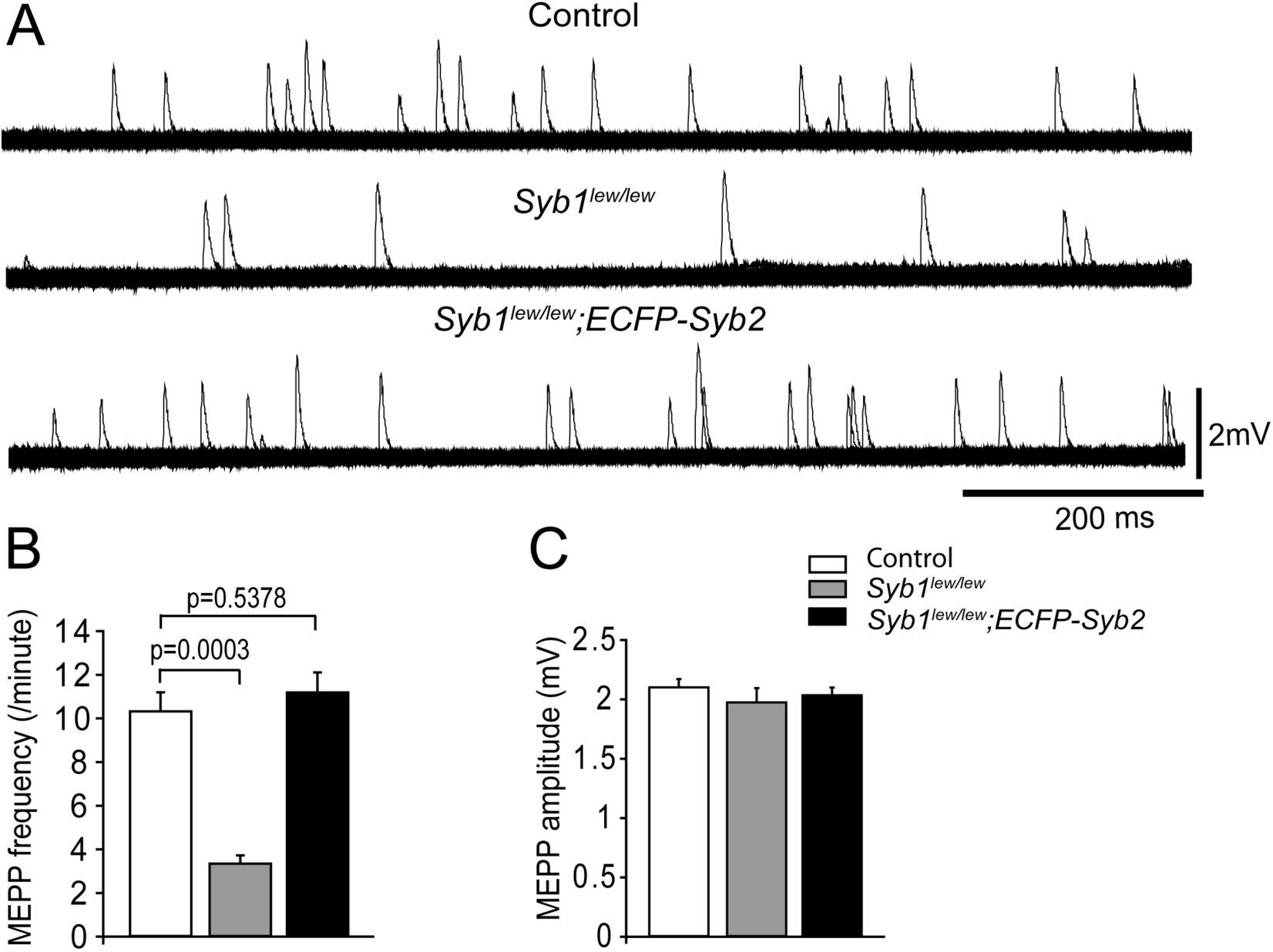
Transgenic ECFP-Syb2 expression restores spontaneous neurotransmission at the NMJ in *Syb1^lew/lew^*;ECFP-Syb2 mice. **A**: Representative mEPP traces show markedly reduced spontaneous activity in *Syb1^lew/lew^* mice, which is restored in *Syb1^lew/lew^;ECFP-Syb2* mice. Representative mEPP traces recorded from diaphragm muscles of control, *Syb1^lew/lew^*and *Syb1^lew/lew^;ECFP-Syb2* mice at P14. 120 consecutive 1-second sweeps are superimposed. **B**: Quanticifation of mEPP frequency and amplitude. The MEPP frequency is significantly reduced in the *Syb1^lew/lew^* mice (3.29 ± 0.38 per minute), but is restored to the control level in *Syb1^lew/lew^;ECFP-Syb2* mice (control: 10.21 ± 0.86 per min; *Syb1^lew/lew^;ECFP-Syb2*: 11.06 ± 0.91 per minute). mEPP amplitude remains unchanged across genotypes. *Syb1^lew/^*^lew^ and *Syb1^lew/lew^;ECFP-Syb2* mice (control: 2.1 ± 0.09 mV; *Syb1^lew/lew^*: 1.98 ± 0.14 mV; *Syb1^lew/lew^;ECFP-Syb2*: 2.04 ± 0.07 mV). The number of mice and cells analyzed: control: N = 3 mice, n = 35 cells; *ECFP-Syb2:* N = 3, n = 51; *Syb1^lew/lew^;ECFP-Syb2*: N = 3, n = 51.

Evoked release was next examined following nerve stimulation. EPP amplitude was significantly reduced in *Syb1^lew/lew^* mice (14.01 ± 1.42 mV) compared to controls (27.92 ± 1.62 mV). This deficit was fully rescued in *Syb1^lew/lew^;ECFP-Syb2* mice (29.93 ± 1.01 mV, n = 18), which were indistinguishable from controls (Fig. 8A, B). Consistently, quantal content was restored to control levels (control: 14.32 ± 0.64; *Syb1^lew/lew^*: 7.03 ± 0.68; *Syb1^lew/lew^;ECFP-Syb2*: 15.49 ± 0.64) (Fig. 8B). Notably, the increased variability in EPP amplitude observed in *Syb1^lew/lew^*mice was normalized in rescued mice (Fig. 8A).

**Figure 8.**
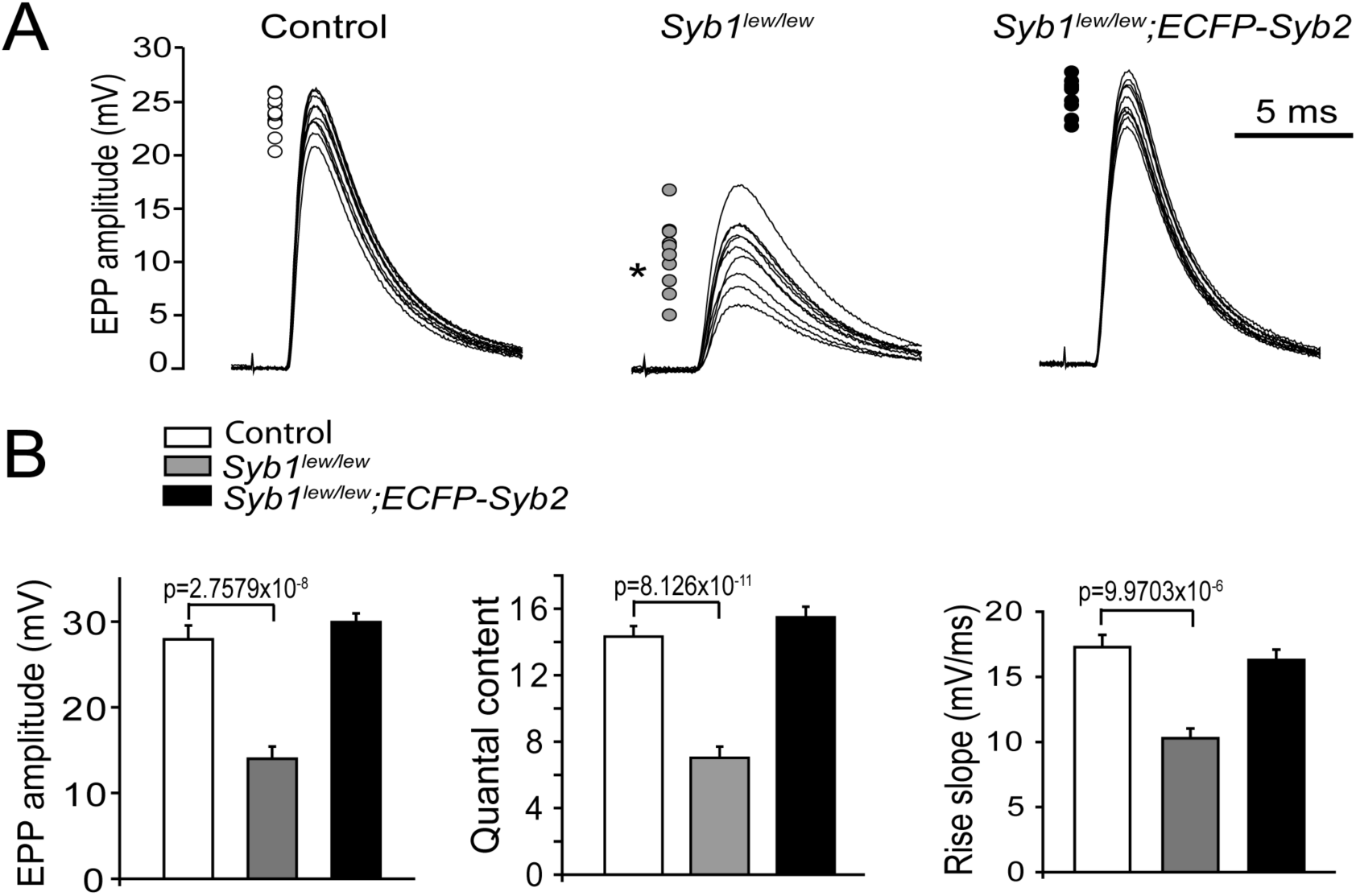
Transgenic ECFP-Syb2 expression rescues evoked neurotransmission at the NMJ in *Syb1^lew/lew^*;ECFP-Syb2 mice. **A**: Evoked EPPs are highly reproducible in control mice, but markedly variable in *Syb1^lew/lew^* mice; ECFP-Syb2 expression restores EPPs to control levels. Ten consecutive EPP traces (evoked by 0.2 Hz stimulation of the phrenic nerves) are superimposed. The peaks of all 10 EPPs are indicated by circles located to the left of the traces. An F-test demonstrated that the variances in *Syb1^lew/lew^* mice were significantly increased (P = 0.0052) compared to those of control mice. The variability and EPP amplitudes in *Syb1^lew/lew^;ECFP-Syb2* mice are restored to the levels of the control mice. **B:** EPP amplitude, quantal content, and rise slope are significantly reduced in *Syb1^lew/lew^* mice but are fully restored to control values in *Syb1^lew/lew^;ECFP-Syb2* mice, indicating complete rescue of evoked synaptic transmission. EPP amplitudes: control, 27.92 ± 1.62 mV; *Syb1^lew/lew^*, 14.01 ± 1.42 mV; *Syb1^lew/lew^;ECFP-Syb2*, 29.93 ± 1.01 mV. Quantal contents: control, 14.32 ± 0.64; *Syb1^lew/lew^*, 7.03 ± 0.68; *Syb1^lew/lew^;ECFP-Syb2*, 15.49 ± 0.64. Rise slopes: control, 15.59 ± 1.24; *Syb1^lew/lew^*, 8.72 ± 0.83; *Syb1^lew/lew^;ECFP-Syb2*, 16.29 ± 0.73. The number of mice and cells analyzed: control: N = 3 mice, n = 21cells, *Syb1^lew/lew^*: N = 3, n = 30; *Syb1^lew/lew^;ECFP-Syb2*: N = 3, n = 27.

We further assessed synaptic short-term plasticity. During 30 Hz train stimulation, *Syb1^lew/lew^* NMJs displayed pronounced facilitation relative to controls. In contrast, *Syb1^lew/lew^; ECFP-Syb2* NMJs exhibited a normal response profile, with modest initial facilitation followed by progressive synaptic depression, comparable to controls (Fig. 9).

**Figure 9.**
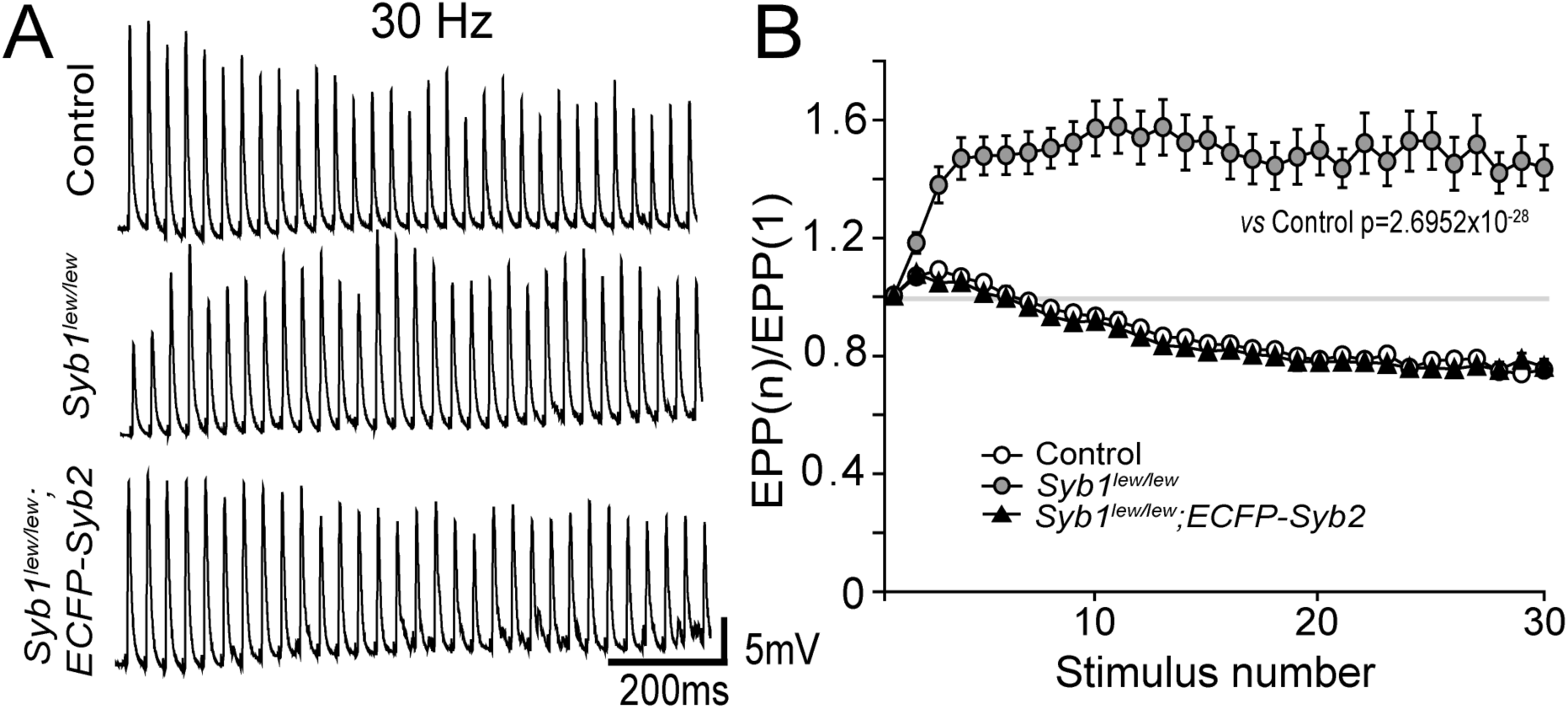
Transgenic ECFP-Syb2 expression restores short-term synaptic plasticity at the NMJ in *Syb1^lew/lew^*;ECFP-Syb2 mice. **A**: Representative EPP responses during 30 Hz train stimulation (1 second) show altered synaptic dynamics in *Syb1^lew/lew^* mice compared with control and rescued mice. Sample EPP traces are shown in response to 30 Hz train stimulation (1 second train). **B**: Quantification of EPP run-down rate calculated by the ratio of EPP amplitude to the first EPP amplitude [EPP(n)/EPP(1)]. The NMJs in *Syb1^lew/lew^* exhibit abnormal short-term plasticity characterized by enhanced facilitation and altered run-down kinetics, whereas the NMJs in *Syb1^lew/lew^;GFP--Syb2* mice display EPP run-down profiles comparable to control, indicating restoration of normal synaptic plasticity. The number of mice and cells analyzed: control: N = 3 mice, n =19 cells; *Syb1^lew/lew^* : N = 3, n = 21; *Syb1^lew/lew^;ECFP-Syb2*: N = 3, n = 18.

Together, these results demonstrate that transgenic *ECFP-Syb2* fully restores synaptic neurotransmission at *Syb1^lew/lew^* NMJs.

## Discussion

We previously showed that loss of SYB1 causes severe neuromuscular transmission defects, including reduced spontaneous and evoked release and impaired Ca²^+^ sensitivity and cooperativity (Liu et al., 2011). In the present study, we demonstrate that neuron-specific expression of Syb2 is sufficient to rescue this lethal phenotype. Introduction of a *Thy1*-driven *ECFP-Syb2* transgene into *Syb1^lew/lew^* mice restores survival, normal growth, and motor function. Electrophysiological analyses further demonstrate complete correction of spontaneous and evoked transmission defects at Syb1-deficient NMJs. Together, these findings demonstrate that Syb2 can functionally substitute for Syb1 at motor nerve terminals *in vivo*.

Syb1/VAMP1 and Syb2/VAMP2 are closely related vesicular SNARE isoforms with high sequence homology but partially distinct and overlapping expression patterns in the nervous system (Elferink et al., 1989; Jacobsson et al., 1998; Li et al., 1996; Raptis et al., 2005; Sudhof et al., 1989). Multiple studies have suggested that specific neuronal populations preferentially utilize one isoform, indicating potential isoform-specialized functions (Jacobsson et al., 1998; Meng et al., 2007; Sherry et al., 2003). For example, in the mouse retina, Syb1 and Syb2 show differential synaptic distribution: Syb2 is broadly expressed at both ribbon and conventional synapses, whereas Syb1 is more restricted and is also found in dendrites, somata, and axons of certain retinal ganglion cells (Sherry et al., 2003).

In our previous studies (Liu et al., 2011; Liu et al., 2019) and the present work, we show that Syb1 and Syb2 are co-expressed at embryonic, juvenile, and adult NMJs. Functionally, Syb2 deficiency causes enhanced spontaneous release, reduced evoked transmission, and perinatal lethality, whereas Syb1 deficiency leads to normal embryonic NMJ transmission but postnatal lethality at ∼3 weeks, suggesting a greater role for Syb2 during early development. In Syb1/Syb2 double-mutant mice, evoked neurotransmission is completely abolished, demonstrating that these two isoforms together account for the majority of synaptobrevin-dependent vesicle fusion at the NMJ (Liu et al., 2019).

In the present study, we show that neuron-specific expression of ECFP-Syb2 fully rescues both spontaneous and evoked neurotransmission in Syb1-deficient NMJs and restores survival and motor function. These findings indicate that ECFP-Syb2 is properly incorporated into the SNARE complex and functions physiologically at motor nerve terminals in the absence of Syb1. Together, our results demonstrate that increasing neuronal Syb2 can compensate for Syb1 loss, supporting the conclusion that Syb1 and Syb2 perform largely overlapping roles in synaptic vesicle fusion at the NMJ.

In addition to Syb1 and Syb2, other synaptobrevin isoforms, including VAMP4 and VAMP7 (also known as tetanus toxin–insensitive VAMP, TI-VAMP), are expressed in neurons, albeit at relatively low levels (Advani et al., 1998; Antonin et al., 2000; Bal et al., 2013; Muzerelle et al., 2003; Raingo et al., 2012; Ramirez et al., 2012; Scheuber et al., 2006). These isoforms appear to serve more specialized functions in membrane trafficking. For example, at central synapses, VAMP4 has been implicated in Ca²^+^-dependent asynchronous release, whereas Syb2 primarily mediates fast synchronous neurotransmitter release (Raingo et al., 2012). Given the structural and functional specialization of the NMJ relative to central synapses, it remains unclear whether VAMP4, VAMP7, or other synaptobrevin isoforms are expressed in motor neurons or are specifically localized to the NMJ. Clarifying their expression and potential contributions will be important for defining the molecular mechanisms that regulate and sustain neurotransmitter release at this peripheral synapse.

Although Syb1 is one of the two major synaptobrevin isoforms in the nervous system, it has also been detected in multiple non-neuronal tissues (Ferlito et al., 2010; Jagadish et al., 1996; Ralston et al., 1994; Rossetto et al., 1996). Syb1 has been detected in rat smooth muscle, heart, pancreas, liver, and kidney (Rossetto et al., 1996), consistent with a broader role in regulated exocytosis. In insulin-responsive tissues such as skeletal muscle and adipocytes, Syb1 and Syb2 assemble with syntaxin-4 and SNAP-25 to mediate fusion of GLUT4-containing vesicles with the plasma membrane (Jagadish et al., 1996). Similarly, Syb1, Syb2, and syntaxin-4 participate in SNARE complexes in atrial myocytes to regulate exocytotic release of atrial natriuretic peptide (Ferlito et al., 2010; Hinners et al., 2003; Peters et al., 2006). Despite this widespread expression, our data demonstrate that *Syb1^lew/lew^* mice can be fully rescued by neuron-specific expression of ECFP-Syb2. This finding indicates that the essential requirement for Syb1 *in vivo* resides within the nervous system, whereas its non-neuronal functions are not required for postnatal survival in mice.

Pathogenic variants in *Syb1* have been associated with a spectrum of human neuromuscular disorders. In particular, SYB1 has been identified as a presynaptic protein implicated in congenital myasthenic syndrome (CMS), with homozygous or compound heterozygous mutations reported in patients with autosomal recessive CMS (Polavarapu et al., 2021; Salpietro et al., 2017; Shen et al., 2017; Yildirim et al., 2024). In addition, heterozygous mutations in Syb1 have been linked to dominantly inherited hereditary spastic ataxia (Bourassa et al., 2012; Grewal et al., 2004; Meijer et al., 2002). In this context, our findings provide insight into the functional consequences of SYB1 deficiency *in vivo* and may inform future efforts aimed at developing therapeutic strategies for SYB1-related neurological disorders.

## Acknowledgments

This work was supported by grants from NIH/NINDS (R01 NS055028), the Edward Mallinckrodt, Jr. Foundation, and the Cain Foundation in Medical Research.

